# Phylogenetic Insights into Antifungal Susceptibility: Comparing Environmental and Clinical *Scedosporium* and *Lomentospora* Isolates from Taiwan

**DOI:** 10.1101/2025.11.08.687386

**Authors:** Shang-Yi Lin, Yi-Ting Tseng, Khaled Abdrabo El-Sayid Abdrabo, Hsin-Mao Wu, Pei-Lun Sun, Chi-Jung Wu, Passanesh Sukphopetch, Sharon C.-A. Chen, Yin-Tse Huang

## Abstract

**Objectives:** *Scedosporium* and *Lomentospora* species are intrinsically resistant molds increasingly implicated in invasive infections with poor outcomes. Olorofim, a novel orotomide antifungal, has demonstrated potent activity against various molds. This study evaluated the antifungal susceptibility of clinical and environmental isolates, focusing on olorofim, and examined these data in a phylogenetic context.

**Methods:** A total of 216 isolates (*Scedosporium* spp., n=205; *L. prolificans*, n=11) were collected from clinical and environmental sources across Taiwan. Species identification was confirmed by sequencing of the ITS and β-tubulin regions, followed by maximum-likelihood phylogenetic analysis. Antifungal susceptibility was determined using CLSI M38-A3 method. Cross-resistance was evaluated using correlation analyses. Phylogenetic signal, Mantel tests, and variance partitioning were applied to assess the evolutionary structuring of minimum inhibitory concentrations (MICs).

**Results:** *Scedosporium apiospermum* (46.3%) and *S. boydii* (15.7%) were the predominant species, exhibiting comparable triazole susceptibility profiles (MIC_90_: voriconazole 1-2 mg/L, posaconazole 4 mg/L, and isavuconazole 8 mg/L). *L. prolificans* displayed pan-resistance to all agents except olorofim. Among triazoles, voriconazole yielded the lowest MICs for most *Scedosporium* isolates. Olorofim exhibited the lowest MICs across *Scedosporium* spp. and *L. prolificans* (all MIC_90_ ≤0.12 mg/L). Rare taxa (*S. marinum*, *S. haikouense*, *S. multisporum*, *S. minutisporum*, *S.* sp. nov.1) were also identified, exhibiting variable triazole MICs but consistently low olorofim MICs. The association between antifungal susceptibility and species relatedness varied by drug: moderate for posaconazole, amphotericin B, and voriconazole; intermediate for olorofim; weak for isavuconazole; and absent for itraconazole. Most MIC variability (72.5–89.2%) occurred within rather than between species.

**Conclusions:** Antifungal susceptibility among *Scedosporium* spp. and *L*. *prolificans* was drug dependent. Olorofim demonstrated consistently low MICs across species, whereas amphotericin B and triazoles showed variable, species-specific activity. These findings highlight olorofim as a promising therapeutic option and emphasize the need for target gene-based investigations to elucidate the evolutionary basis of antifungal resistance.

## Introduction

Infections caused by rare molds are increasingly recognized, particularly among immunocompromised hosts, and the global rise in antifungal resistance has become a major public health concern [1]. *Scedosporium* and *Lomentospora* species are of particular significance due to their intrinsic resistance to multiple antifungal classes [2], a characteristic that recently led to their inclusion in the World Health Organization (WHO) Fungal Priority Pathogen List [3]. These environmental molds can cause a wide range of diseases, from localized infections to life-threatening invasive disease, in both immunocompromised and immunocompetent individuals, with reported mortality rates remaining exceedingly high (46–100%) [2, 3].

*Scedosporium* spp. *and L. prolificans* have emerged as clinically significant filamentous fungi with distinct epidemiological and virulence characteristics [4, 5]. Their adaptability to human-modified environments, including recreational areas, polluted soils, and contaminated water, creates extensive ecological reservoirs that facilitate opportunistic infection [6]. Recent environmental surveys in Taiwan confirmed their widespread prevalence in soil, particularly in areas with substantial human activity [7]. Such ecological insights are essential for identifying potential infection sources and predicting outbreak risks [8, 9].

The *in vitro* antifungal susceptibility of *Scedosporium* spp. and *Lomentospora prolificans* varies considerably. While *Scedosporium* spp. generally exhibit partial susceptibility to triazoles, *L. prolificans* remains resistant to nearly all currently available antifungal agents [2, 3]. This high level of intrinsic resistance poses significant challenges for clinical management. Olorofim, the first-in-class orotomide antifungal, disrupts the fungal *de novo* pyrimidine biosynthetic pathway through selective inhibition of fungal dihydroorotate dehydrogenase (DHODH), resulting in deleterious effects on RNA and DNA synthesis [10]. It has demonstrated potent *in vitro* activity against a broad range of clinically relevant molds [11–13]. Recent studies from the United States, Australia, and Spain further support its strong activity against *Scedosporium* spp. and *L. prolificans* [11, 13–15]. However, because antifungal susceptibility patterns can vary across geographic regions [16, 17], region-specific studies are essential to guide appropriate therapeutic strategies.

Despite their clinical relevance, substantial knowledge gaps remain regarding the antifungal susceptibility of *Scedosporium* and *Lomentospora* in Taiwan, particularly in relation to environmental reservoirs and evolutionary factors. This study investigated the antifungal susceptibility of *Scedosporium* and *Lomentospora* isolates from Taiwan, comparing clinical and environmental strains, integrating results with phylogenetic analyses, and evaluating the activity of novel antifungal agents such as olorofim.

## Materials and Methods

### Isolate Selection, Identification, and Phylogenetic Analysis

A total of 216 isolates were analyzed, comprising 205 *Scedosporium* spp. and 11 *L. prolificans* isolates obtained from clinical and environmental sources in Taiwan. Environmental sampling yielded 168 isolates from soil at 50 locations across Taiwan between 2014 and 2024 (**Figure S1**).

Sampling sites represented varying levels of human activity, including urban areas (disturbed sites), hospital environments (defined as soil collected from hygiene-sensitive areas within hospital grounds), and natural sites (less-disturbed regions). Detailed sampling procedures have been described previously [7]. A total of 48 clinical isolates were collected from three medical centers in Taiwan between 1 January 2012 and 31 July 2022 (**Table S1, Figure S1**). The study was reviewed and approved by the Institutional Review Board of Kaohsiung Medical University Hospital (IRB no. KMUHIRB-E(I)-20250290).

Species identification was confirmed by polymerase chain reaction (PCR) amplification and sequence analysis of the internal transcribed spacer (ITS) region of the ribosomal DNA (rDNA) and part of the β-tubulin (BT2) gene, following previously established protocols [7]. ITS and BT2 sequences were aligned separately using Multiple Alignment using Fast Fourier Transform (MAFFT) with default parameters, trimmed using ClipKIT [18], and concatenated for further analysis. The optimal nucleotide substitution models for each gene region were determined using ModelTest-NG. Maximum likelihood (ML) phylogenetic trees were generated in IQ-TREE2 with 1,000 bootstrap replicates, applying partition parameters recommended by ModelTest-NG. The resulting phylogenetic tree was visualized using Interactive Tree of Life (iTOL) v6.

### Antifungal Susceptibility Testing

Antifungal susceptibility testing was conducted in accordance with the Clinical and Laboratory Standards Institute (CLSI) M38-A3 guidelines [19]. The minimum inhibitory concentration (MIC) was defined as the lowest concentration that completely inhibited visible growth compared with the drug-free control well. Final concentration ranges for each antifungal were as follows: olorofim, 0.001 to 1 mg/L; amphotericin B and itraconazole, 0.015-16 mg/L; and voriconazole, posaconazole, and isavuconazole, 0.008-8 mg/L. Olorofim was provided by F2G Ltd. (Cheshire, United Kingdom), and all other antifungal agents were obtained from Sigma (St. Louis, MO). Quality control was ensured by testing the following reference strains daily: *Candida parapsilosis* (ATCC 22019), *Aspergillus fumigatus* (ATCC MYA-3626), *Aspergillus flavus* (ATCC 204304), and *Scedosporium apiospermum* (ATCC MYA-3635).

### Antifungal Cross-Resistance Analysis

Cross-resistance patterns among antifungal agents were analyzed in R (version 4.3.1) using the dplyr [20] and corrplot [21] packages. MIC data for six antifungal agents were evaluated. Because MIC distributions were non-normal, Spearman’s rank correlation coefficients (ρ) were calculated for all pairwise drug comparisons using the cor() function with pairwise complete observations to account for missing data. Statistical significance for each correlation was assessed using the cor.test() with the Spearman method. Correlation matrices were visualized with corrplot() function, employing hierarchical clustering to group antifungals with similar resistance patterns.

### Phylogenetic Comparative Analysis of Antifungal Susceptibility Patterns

To determine whether antifungal susceptibility patterns were phylogenetically conserved or randomly distributed across the phylogeny, phylogenetic comparative analyses were performed as illustrated in Figure S2. MIC values were logC-transformed, and zero-length branches in the phylogenetic tree were assigned a minimum length of 1 × 10CC to avoid computational singularities.

Pagel’s lambda (λ) was calculated using the phylosig() function in the phytools package [22] to quantify phylogenetic signal for each antifungal. Lambda values range from 0 (no phylogenetic dependence) to 1 (trait evolution consistent with Brownian motion). Statistical significance was evaluated using likelihood ratio tests comparing the observed λ to a null model (λ = 0), with *p*-values derived from a chi-square distribution. The relationship between phylogenetic distance and antifungal susceptibility was evaluated using the mantel() function in the vegan package [23]. Pairwise distance matrices of MIC values were compared with phylogenetic distance matrices using Pearson correlation with 999 permutations to assess statistical significance. Analyses were restricted to antifungal agents with sufficient data (>10 isolates with non-missing values). To partition the relative contributions of across-species versus within-species variation, hierarchical clustering was performed on the phylogenetic distance matrix using the cophenetic.phylo() function in the ape package [24] with complete linkage. Analysis of variance (ANOVA) was used to partition total MIC variance into between-species and within-species components, and the proportion of variance explained by each component was calculated for each antifungal agent.

## Results

### Distribution of Fungal Isolates and Antifungal Susceptibility Data

Antifungal susceptibility testing for 216 *Scedosporium* and *L. prolificans* isolates is summarized in **Table 1**. *S. apiospermum* (n=100, 46.3%) and *S. boydii* (n=34, 15.7%), the predominant species, exhibited similar triazole susceptibility profiles, with voriconazole showing the lowest MICs (MIC_90_: voriconazole 1-2 mg/L, posaconazole 4 mg/L, and isavuconazole 8 mg/L). For *Scedosporium* sp. nov.1, posaconazole yielded lower MICs than voriconazole, whereas relatively high posaconazole MICs were observed for *S. marinum*, *S. fusoideum*, and *S. ellipsoideum*. Amphotericin B, itraconazole and isavuconazole demonstrated limited activity against *S. aurantiacum*, while voriconazole retained activity comparable to that against *S. apiospermum*. Several rarely described species, such as *S. marinum*, *S. haikouense*, *S. fusoideum*, and *S. sphaerospermum*, displayed high MICs across triazoles and amphotericin B. Across all species, olorofim demonstrated potent *in vitro* activity, with MICs ≤0.12 mg/L, including *L. prolificans*.(**Table 1**)

**Table 1.**
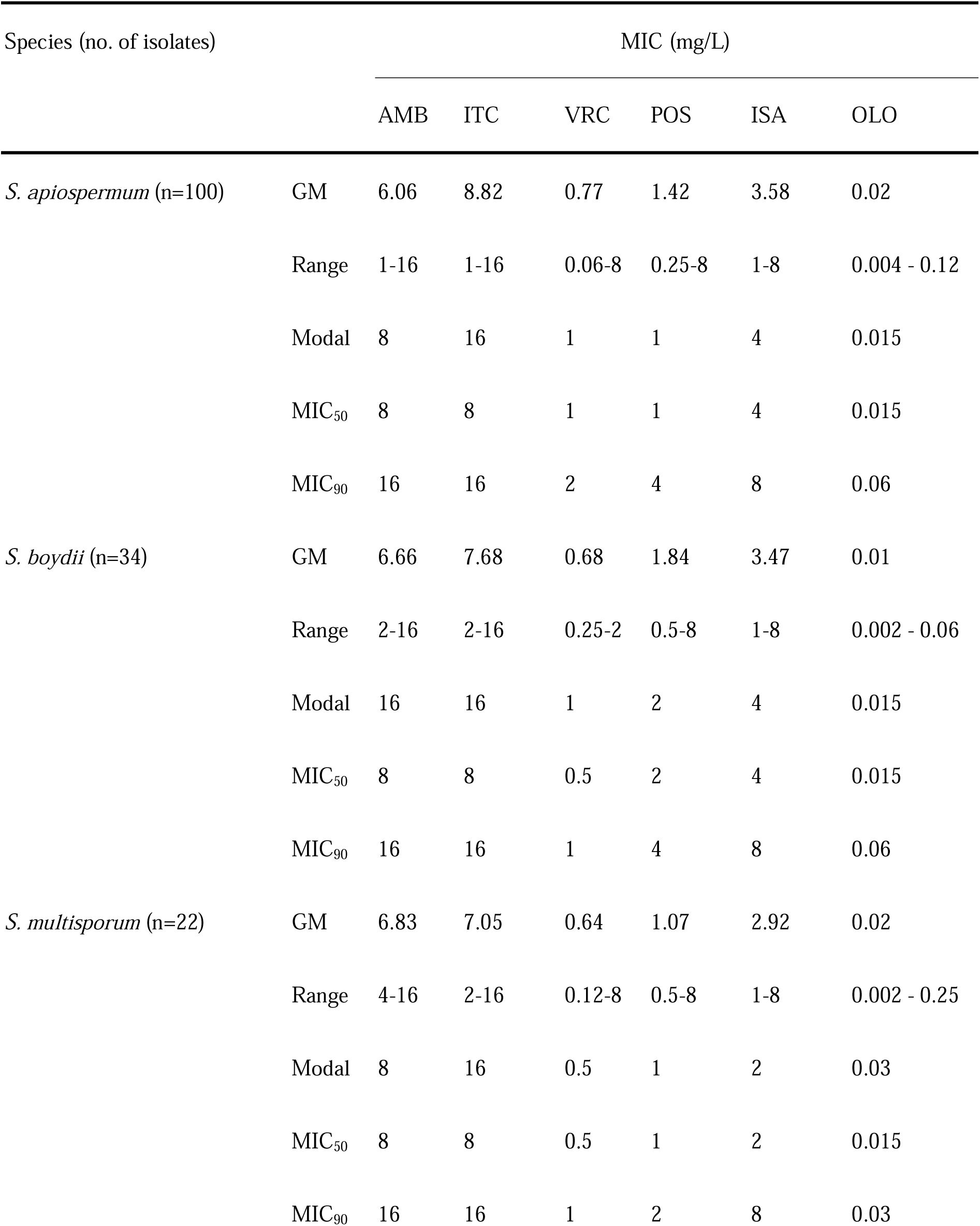
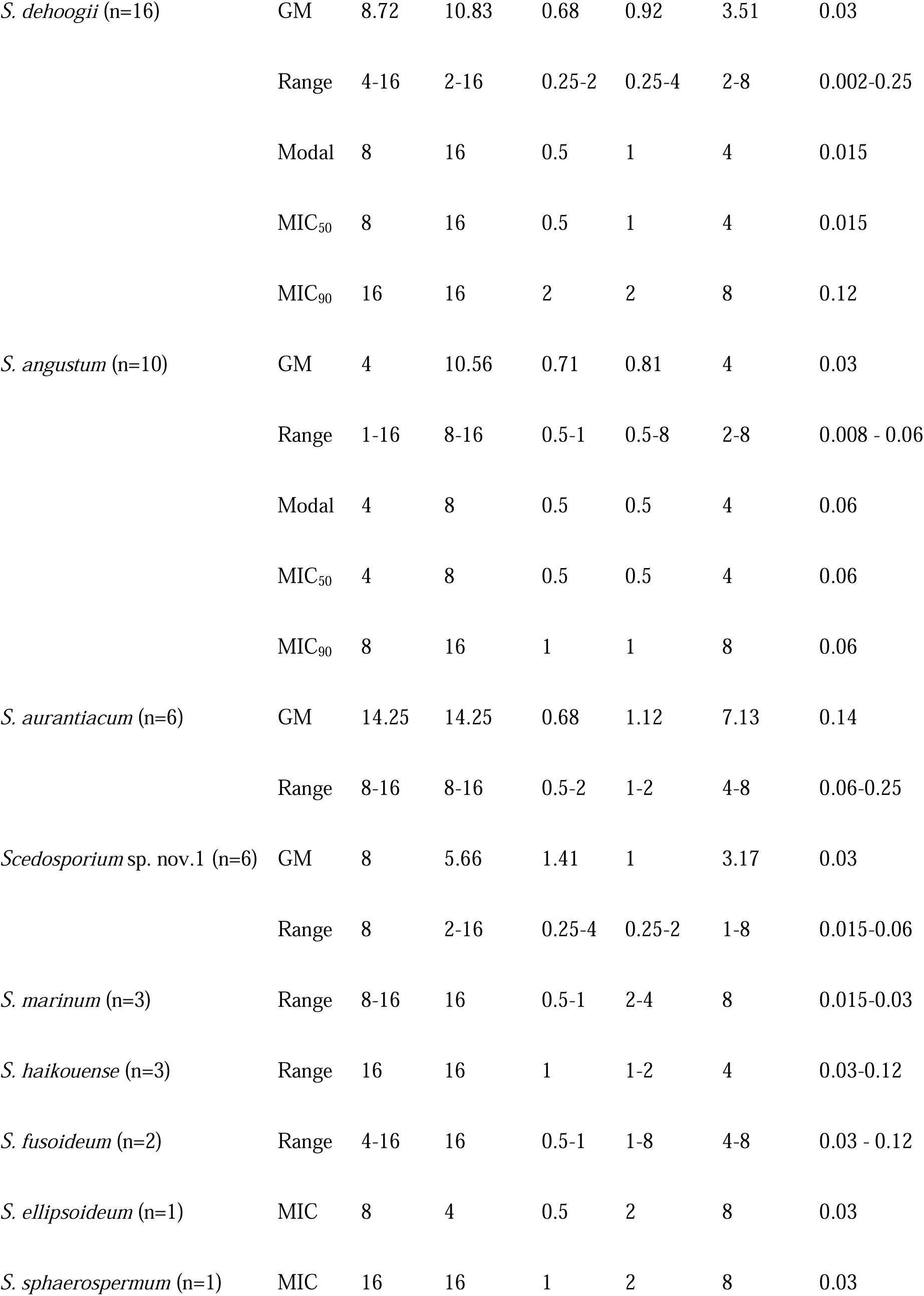
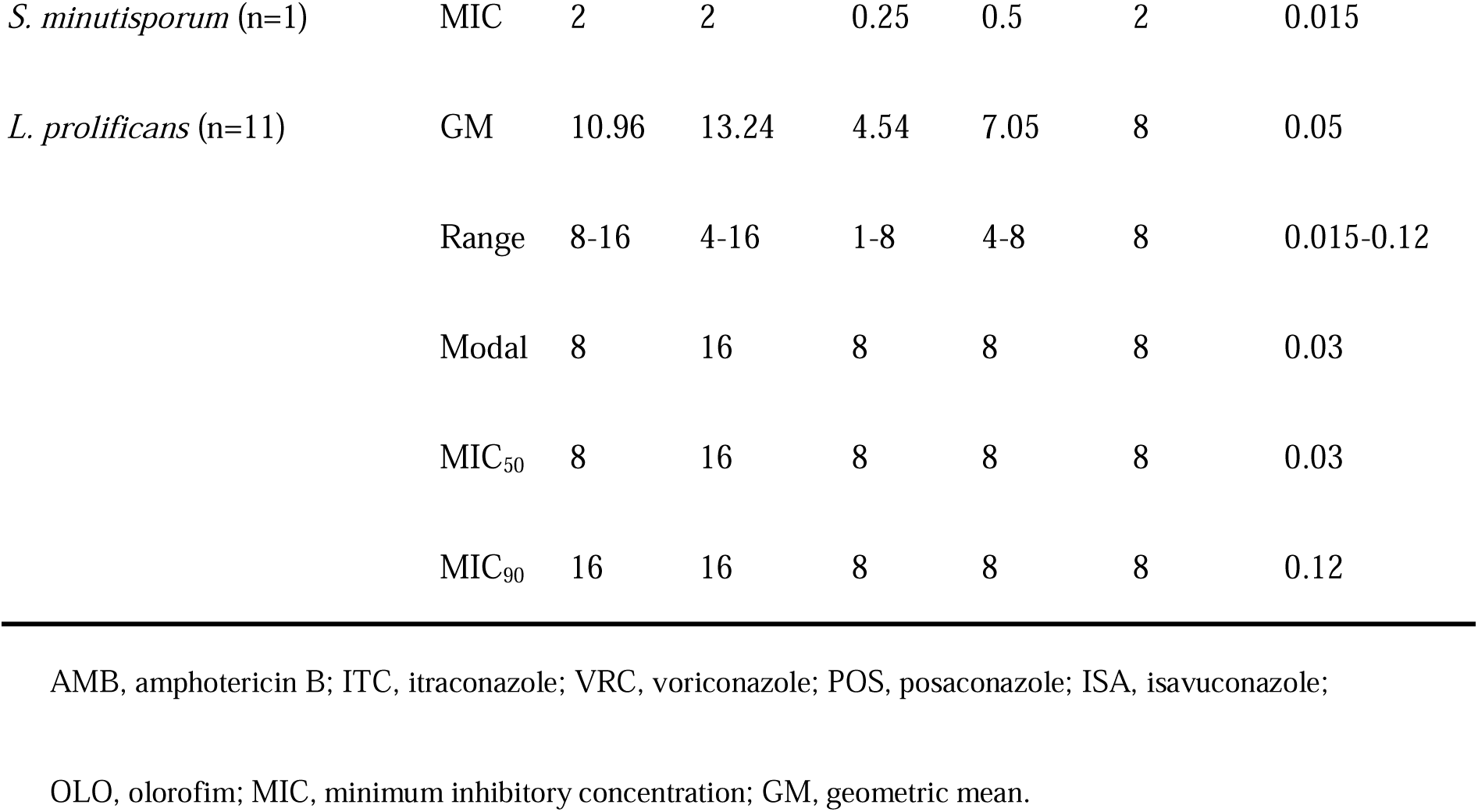
Minimum inhibitory concentration (MIC) range, modal, geometric mean, MIC_50_, and MIC_90_ of *Scedosporium* and *Lomentospora* species (n=216)

| Species (no. of isolates) |  | MIC (mg/L) |  |  |  |  |  |
| --- | --- | --- | --- | --- | --- | --- | --- |
|  |  | AMB | ITC | VRC | POS | ISA | OLO |
| *S. apiospermum* (n=100) | GM | 6.06 | 8.82 | 0.77 | 1.42 | 3.58 | 0.02 |
|  | Range | 1-16 | 1-16 | 0.06-8 | 0.25-8 | 1-8 | 0.004 - 0.12 |
|  | Modal | 8 | 16 | 1 | 1 | 4 | 0.015 |
|  | MIC<sub>50</sub> | 8 | 8 | 1 | 1 | 4 | 0.015 |
|  | MIC<sub>90</sub> | 16 | 16 | 2 | 4 | 8 | 0.06 |
| *S. boydii* (n=34) | GM | 6.66 | 7.68 | 0.68 | 1.84 | 3.47 | 0.01 |
|  | Range | 2-16 | 2-16 | 0.25-2 | 0.5-8 | 1-8 | 0.002 - 0.06 |
|  | Modal | 16 | 16 | 1 | 2 | 4 | 0.015 |
|  | MIC<sub>50</sub> | 8 | 8 | 0.5 | 2 | 4 | 0.015 |
|  | MIC<sub>90</sub> | 16 | 16 | 1 | 4 | 8 | 0.06 |
| *S. multisorum* (n=22) | GM | 6.83 | 7.05 | 0.64 | 1.07 | 2.92 | 0.02 |
|  | Range | 4-16 | 2-16 | 0.12-8 | 0.5-8 | 1-8 | 0.002 - 0.25 |
|  | Modal | 8 | 16 | 0.5 | 1 | 2 | 0.03 |
|  | MIC<sub>50</sub> | 8 | 8 | 0.5 | 1 | 2 | 0.015 |
|  | MIC<sub>90</sub> | 16 | 16 | 1 | 2 | 8 | 0.03 |
| *S. dehoogii* (n=16) | GM | 8.72 | 10.83 | 0.68 | 0.92 | 3.51 | 0.03 |
|  | Range | 4-16 | 2-16 | 0.25-2 | 0.25-4 | 2-8 | 0.002-0.25 |
|  | Modal | 8 | 16 | 0.5 | 1 | 4 | 0.015 |
|  | MIC<sub>50</sub> | 8 | 16 | 0.5 | 1 | 4 | 0.015 |
|  | MIC<sub>90</sub> | 16 | 16 | 2 | 2 | 8 | 0.12 |
| *S. angustum* (n=10) | GM | 4 | 10.56 | 0.71 | 0.81 | 4 | 0.03 |
|  | Range | 1-16 | 8-16 | 0.5-1 | 0.5-8 | 2-8 | 0.008 - 0.06 |
|  | Modal | 4 | 8 | 0.5 | 0.5 | 4 | 0.06 |
|  | MIC<sub>50</sub> | 4 | 8 | 0.5 | 0.5 | 4 | 0.06 |
|  | MIC<sub>90</sub> | 8 | 16 | 1 | 1 | 8 | 0.06 |
| *S. aurantiacum* (n=6) | GM | 14.25 | 14.25 | 0.68 | 1.12 | 7.13 | 0.14 |
|  | Range | 8-16 | 8-16 | 0.5-2 | 1-2 | 4-8 | 0.06-0.25 |
| *Scedosporium* sp. nov.1 (n=6) | GM | 8 | 5.66 | 1.41 | 1 | 3.17 | 0.03 |
|  | Range | 8 | 2-16 | 0.25-4 | 0.25-2 | 1-8 | 0.015-0.06 |
| *S. marinum* (n=3) | Range | 8-16 | 16 | 0.5-1 | 2-4 | 8 | 0.015-0.03 |
| *S. haikouense* (n=3) | Range | 16 | 16 | 1 | 1-2 | 4 | 0.03-0.12 |
| *S. fusoides* (n=2) | Range | 4-16 | 16 | 0.5-1 | 1-8 | 4-8 | 0.03 - 0.12 |
| *S. ellipsoideum* (n=1) | MIC | 8 | 4 | 0.5 | 2 | 8 | 0.03 |
| *S. sphaerospermum* (n=1) | MIC | 16 | 16 | 1 | 2 | 8 | 0.03 |
| *S. minutisporum* (n=1) | MIC | 2 | 2 | 0.25 | 0.5 | 2 | 0.015 |
| *L. prolificans* (n=11) | GM | 10.96 | 13.24 | 4.54 | 7.05 | 8 | 0.05 |
|  | Range | 8-16 | 4-16 | 1-8 | 4-8 | 8 | 0.015-0.12 |
|  | Modal | 8 | 16 | 8 | 8 | 8 | 0.03 |
|  | MIC<sub>50</sub> | 8 | 16 | 8 | 8 | 8 | 0.03 |
|  | MIC<sub>90</sub> | 16 | 16 | 8 | 8 | 8 | 0.12 |

AMB, amphotericin B; ITC, itraconazole; VRC, voriconazole; POS, posaconazole; ISA, isavuconazole;
OLO, olorofim; MIC, minimum inhibitory concentration; GM, geometric mean.

Species represented by more than 10 isolates exhibited consistent across-species variation (**Figure 1**). Voriconazole showed the lowest MICs overall but limited activity against *L. prolificans*. Posaconazole displayed intermediate activity with broader MIC distributions (0.25–8 mg/L), while amphotericin B and itraconazole demonstrated poor activity (MIC_90_ ≥8 mg/L). As expected, *L. prolificans* exhibited a multidrug-resistant phenotype, with uniformly high MICs for all triazoles and amphotericin B but consistently low MICs for olorofim.

**Figure 1.**
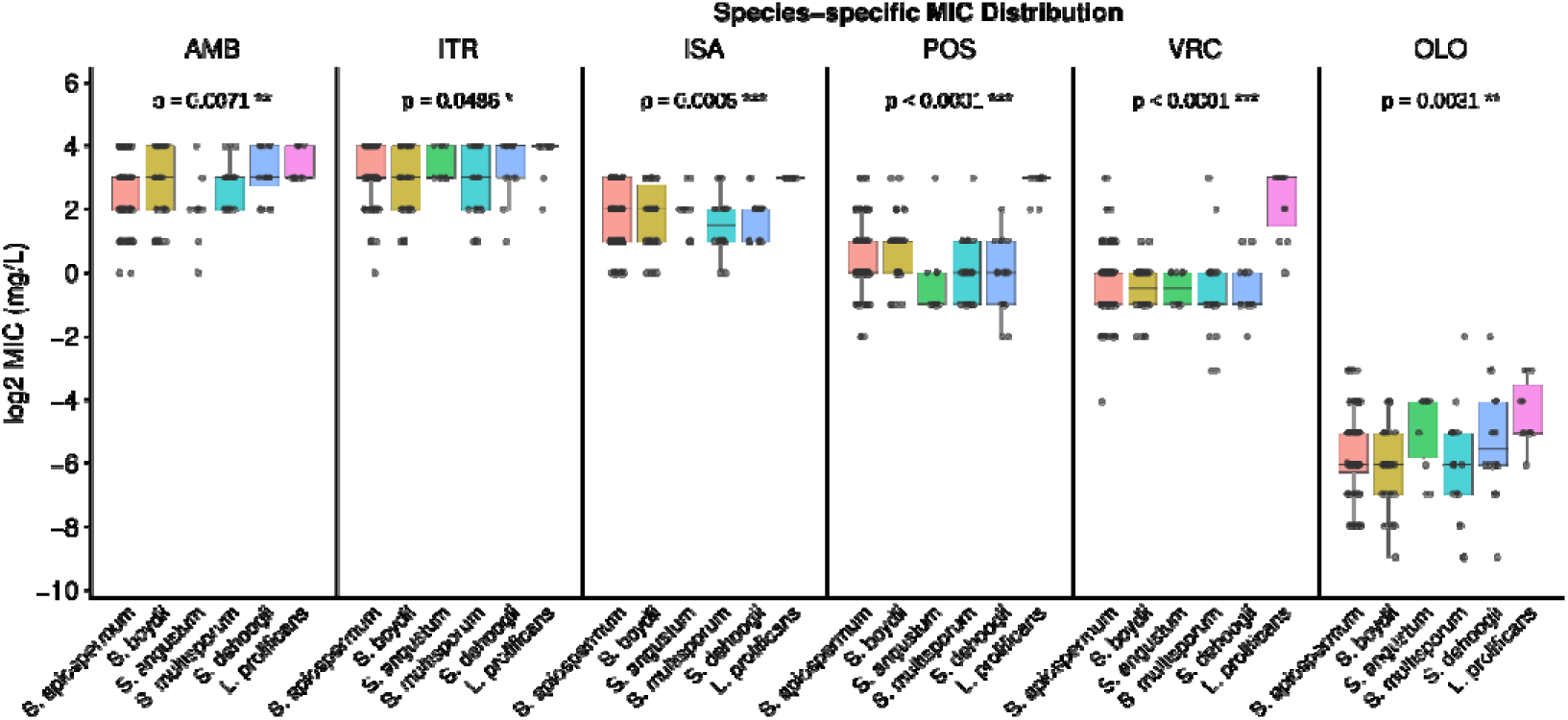
Species-specific minimum inhibitory concentration distributions of *Scedosporium* and *Lomentospora*. Minimum inhibitory concentrations (MICs, mg/L) of olorofim, posaconazole, voriconazole, isavuconazole, amphotericin B, fluconazole, and itraconazole are shown for the most common species: *S. apiospermum* (n=100), *S. boydii* (n=34), *S. angustum* (n=10), *S. multisporum* (n=22), *S. dehoogii* (n=16), and *L. prolificans* (n=11). Each point represents an individual isolate; horizontal lines indicate modal values. MICs are presented on a log2 scale. AMB, amphotericin B; ISA, isavuconazole; ITR, itraconazole; OLO, olorofim; POS, posaconazole; VRC, voriconazole.

Comparison by isolate source (clinical, n=48; hospital environment, n=15; environmental, n=153) revealed that clinical isolates exhibited significantly lower MICs for itraconazole, isavuconazole, and voriconazole than environmental or hospital isolates (all *p* < 0.05). In contrast, MICs for amphotericin B, posaconazole, and olorofim did not differ significantly among sources (**Figure 2**).

**Figure 2.**
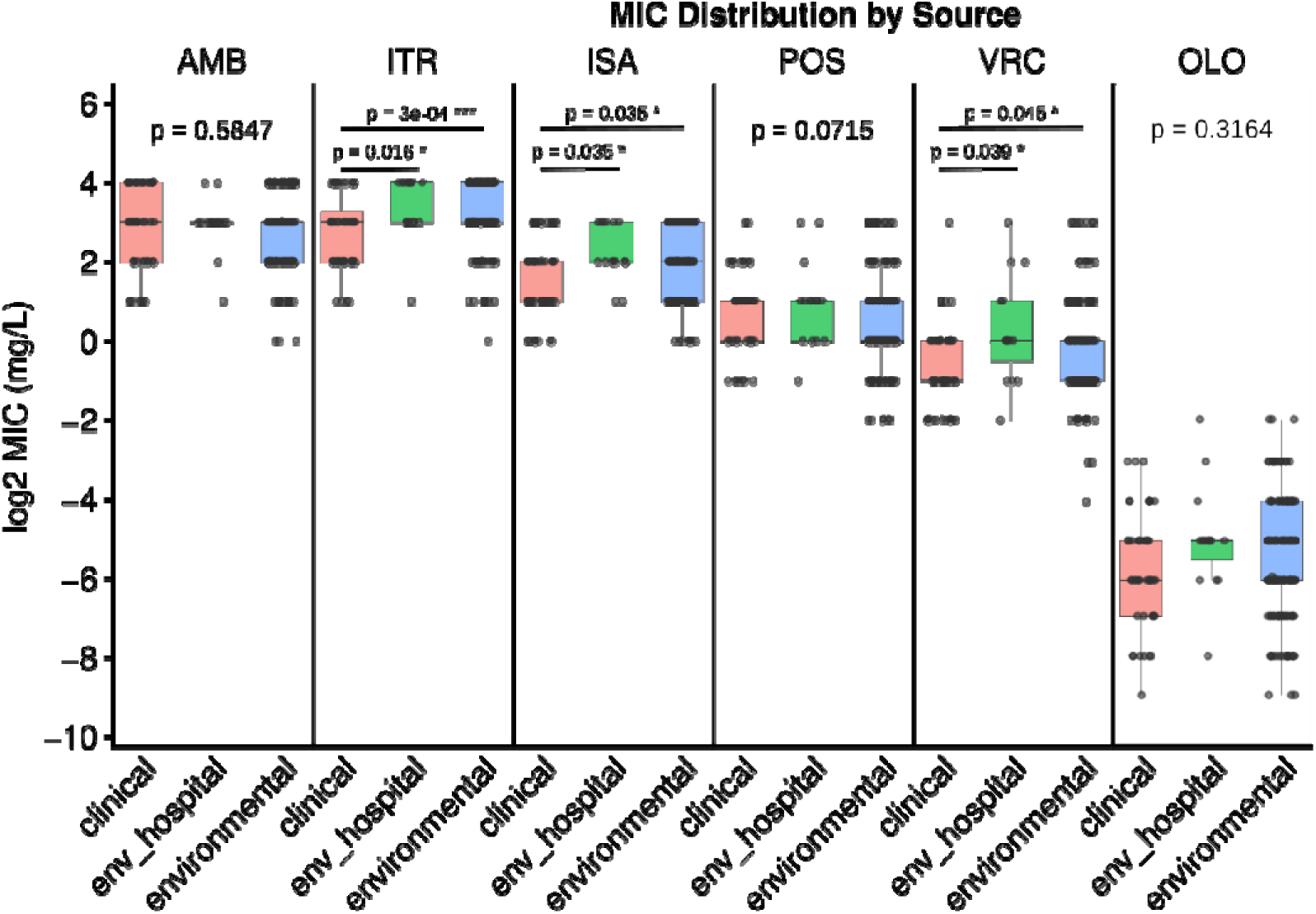
Comparison of minimum inhibitory concentrations between clinical (n=48), hospital environment (n=15), and environmental isolates (n=153) for eight antifungal agents. Each point represents an individual isolate; horizontal lines indicate modal values. AMB, amphotericin B; ISA, isavuconazole; ITR, itraconazole; OLO, olorofim; POS, posaconazole; VRC, voriconazole.

### Antifungal Cross-Resistance Analysis

All antifungal pairs demonstrated positive correlations (ρ = 0.20–0.53), all of which were statistically significant (*p* < 0.05) (**Figure 3**). Among triazoles, isavuconazole and voriconazole demonstrated a moderate correlation (ρ = 0.53, *p* < 0.001). Olorofim showed low correlations with other antifungal agents (ρ = 0.20 to 0.30).

**Figure 3.**
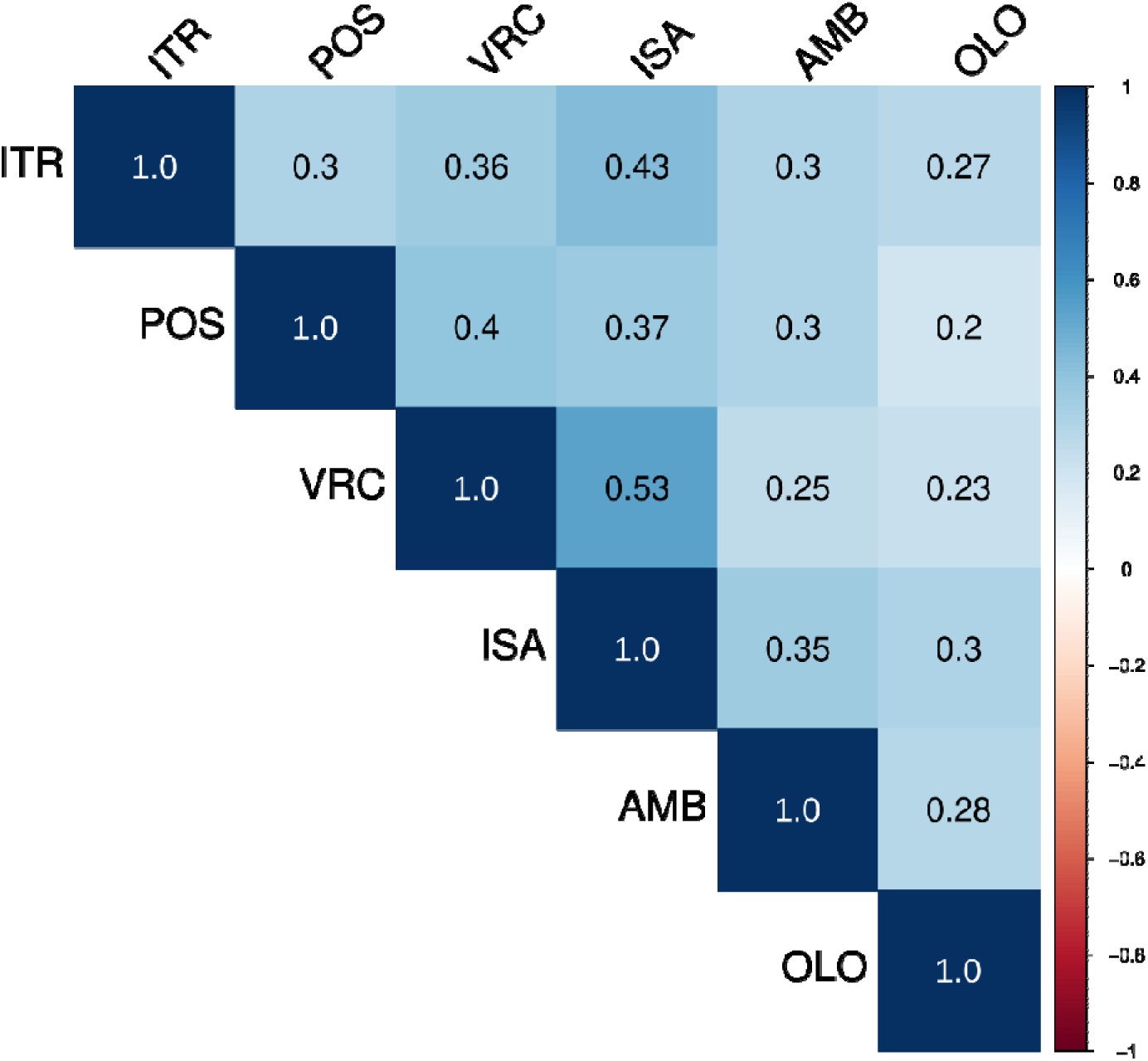
Antifungal cross-resistance analysis. Pairwise correlations between MICs of six antifungal agents are shown. Color intensity represents the strength and direction of Pearson’s correlation coefficient, with blue indicating negative and red indicating positive correlations. AMB, amphotericin B; ISA, isavuconazole; ITR, itraconazole; OLO, olorofim; POS, posaconazole; VRC, voriconazole.

### Phylogenetic Comparative Analysis of Antifungal Susceptibility Patterns

Hierarchical clustering analysis revealed moderate species-dependent patterns in antifungal susceptibility profiles, with no consistent association with isolate source (Figure 4). *L. prolificans* isolates displayed relatively consistent MIC profiles and clustered in specific regions of the dendrogram, whereas other species including *S. apiospermum* and *S. boydii* showed divergent antifungal MIC profiles across isolates, indicating substantial within-species variation.

**Figure 4.**
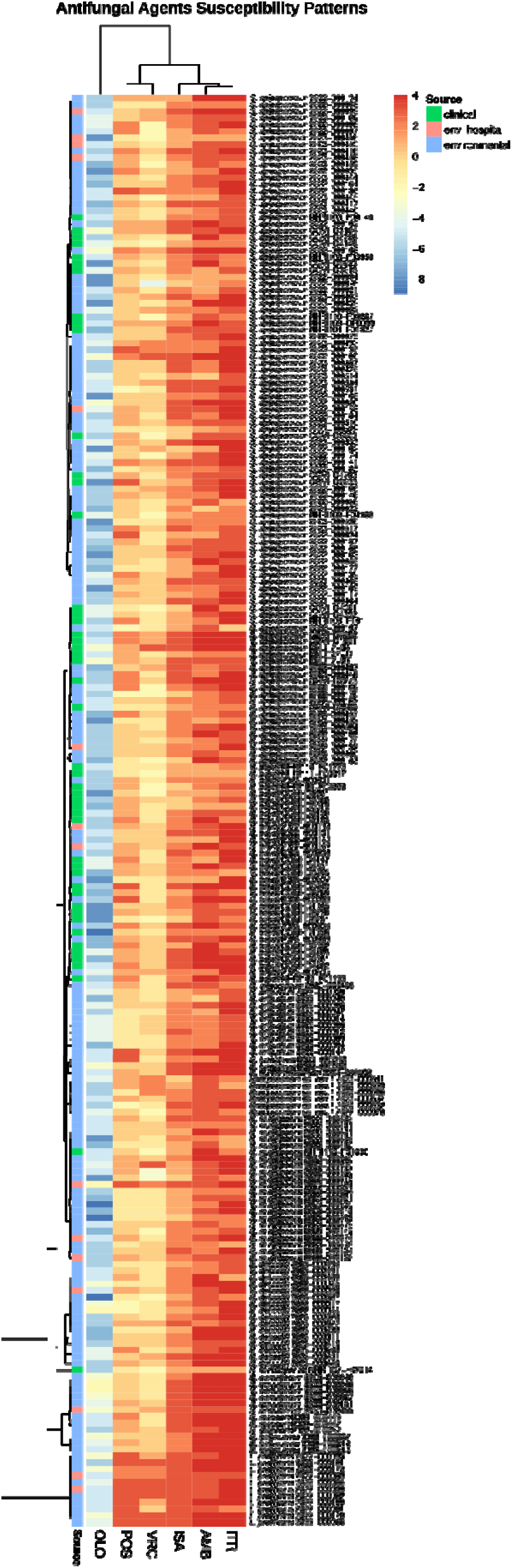
Hierarchical clustering heatmap of scaled log2 minimum inhibitory concentrations for antifungal agents across *Scedosporium* and *Lomentospora* isolates. Rows represent isolates, annotated by species and source (clinical, hospital environmental, or environmental), and columns correspond to antifungal agents. Clustering highlights distinct resistance patterns and associations. AMB, amphotericin B; ISA, isavuconazole; ITR, itraconazole; OLO, olorofim; POS, posaconazole; VRC, voriconazole.

To assess phylogenetic conservation of antifungal susceptibility, three complementary comparative approaches were applied (**Table 2**, **Figure 5**). Phylogenetic signal analysis revealed moderate signals for posaconazole (λ=0.432, p<0.001), amphotericin B (λ=0.405, *p*=0.006), and voriconazole (λ=0.394, p<0.001); an intermediate signal for olorofim (λ=0.357, p=0.001); and a weak signal for isavuconazole (λ=0.230, *p*<0.001). Itraconazole showed no significant signal (λ=0.166, *p*=0.151). Mantel tests demonstrated significant correlations between phylogenetic distance and MIC values for voriconazole (*r*=0.280, *p*=0.001) and posaconazole (*r*=0.239, *p*=0.001), whereas olorofim showed a weaker but still significant correlation (*r*=0.087, *p*=0.021). No significant associations were observed for amphotericin B, itraconazole, or isavuconazole (*p*>0.05). Variance partitioning demonstrated that within-species variation accounted for the majority of MIC heterogeneity (72.5–89.2%). Species effects were more evident for posaconazole and voriconazole (within-species variance: 72.5% and 74.9%, respectively), while itraconazole exhibited the highest within-species variance (89.2%), consistent with its lack of phylogenetic structure.

**Figure 5.**
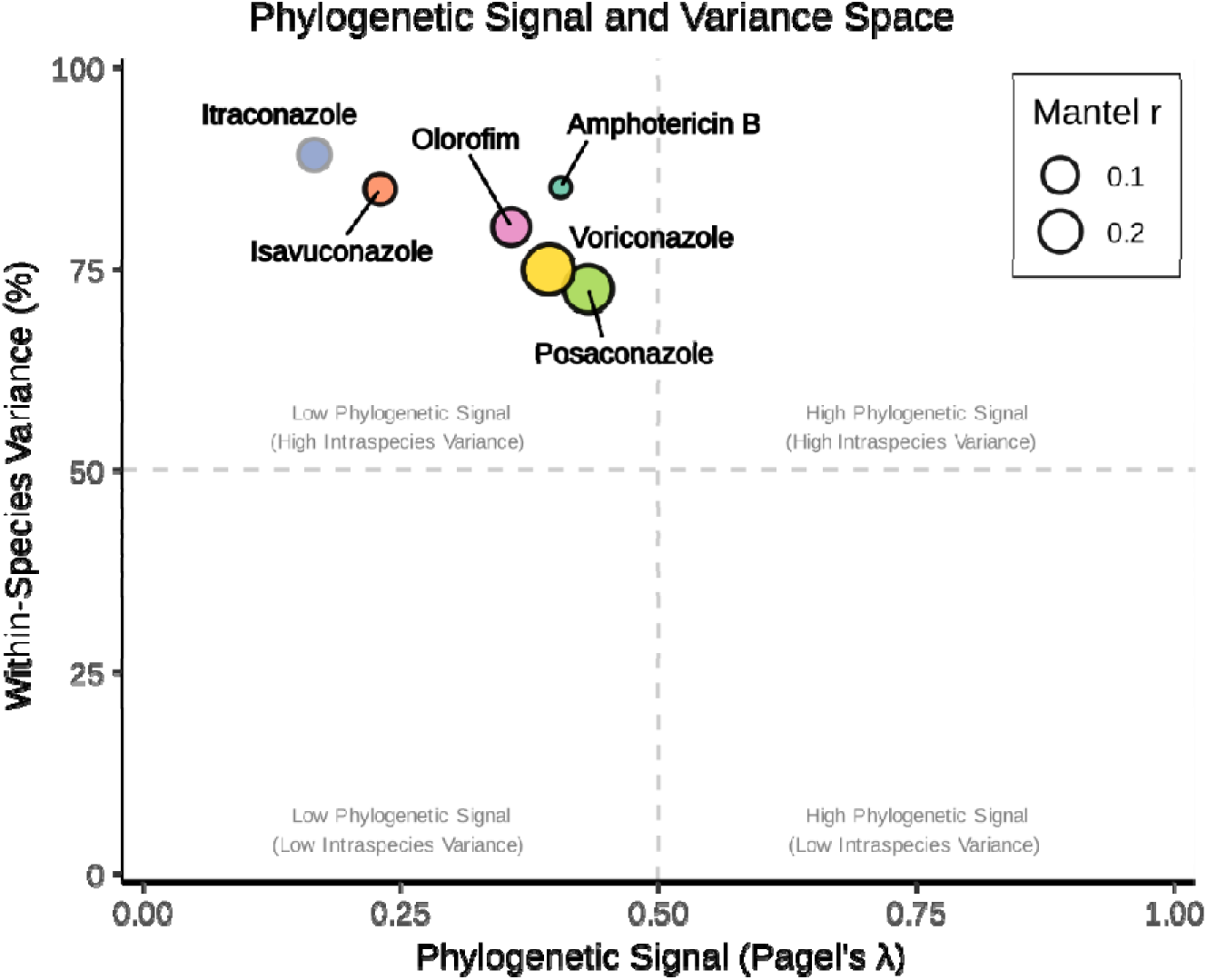
Phylogenetic signal-variance space analysis of antifungal susceptibility patterns. The relationship between phylogenetic signal (Pagel’s λ) and within-species variance (%) for seven antifungal agents. Circle size represents correlation strength between phylogenetic distance and MIC differences (Mantel’s r). Most antifungals show high within-species variance (>70%), with posaconazole and voriconazole displaying the moderate phylogenetic signal, while itraconazole shows minimal phylogenetic signal despite high within-species variance.

**Table 2.**
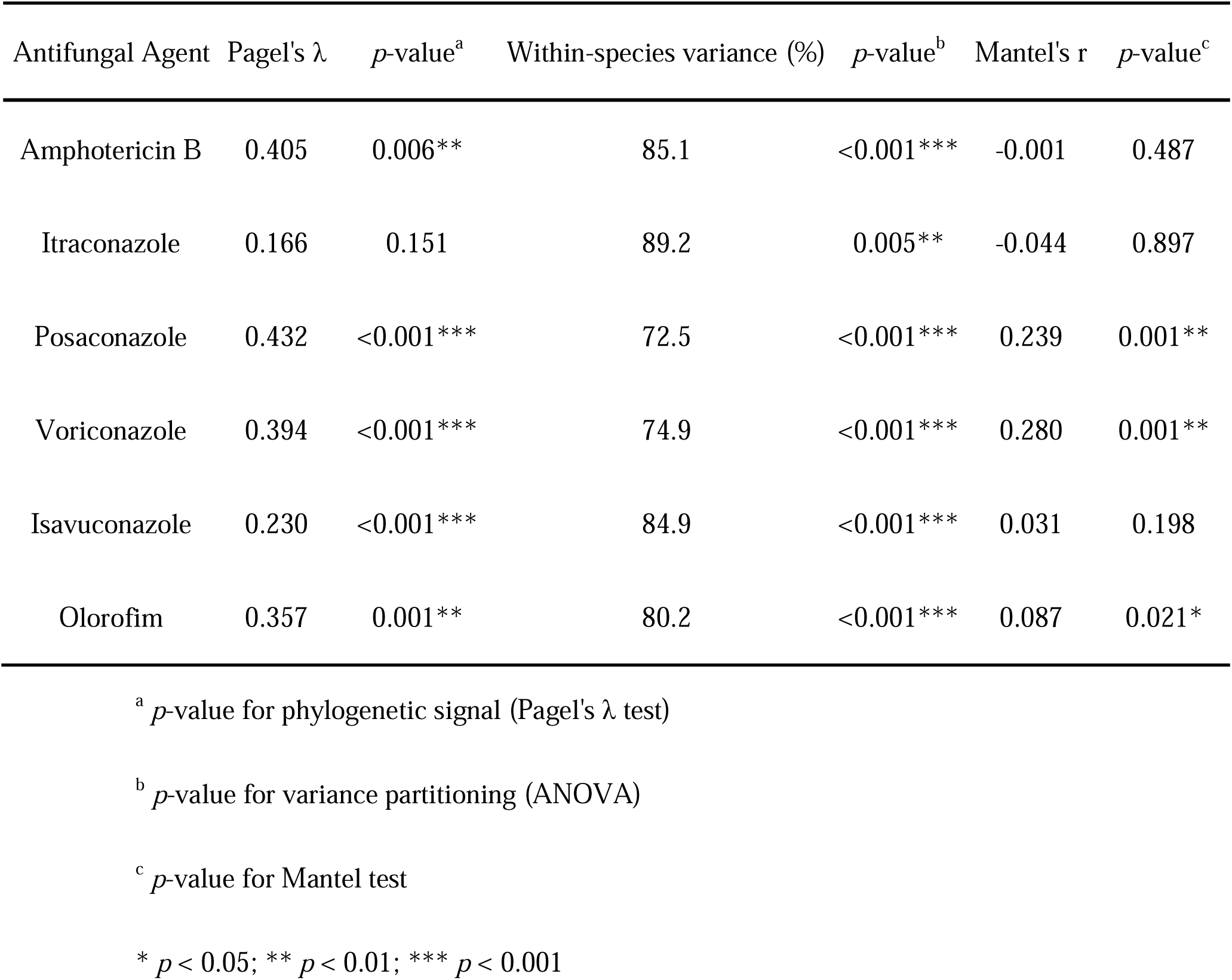
Phylogenetic signal, variance partitioning, and correlation analysis of antifungal susceptibility pattern.

## Discussion

This study provides one of the most comprehensive analysis of antifungal susceptibility among *Scedosporium* species and *L. prolificans*, integrating data from 216 clinical and environmental isolates. We delineated species-specific MIC distributions, evaluated source-related variation, and applied phylogeny-informed methods to evaluate evolutionary structuring. Olorofim demonstrated uniformly low MICs across all taxa, including multidrug-resistant *L. prolificans*, whereas other antifungal agents showed variable, drug-dependent activity patterns. These findings underscore the clinical promise of olorofim and emphasize the importance of integrating molecular identification with susceptibility testing to guide the management of infections caused by emerging resistant molds.

Consistent with previous multicenter surveys, most isolates in our collection belonged to *S. apiospermum* and *S. boydii*, which are the predominant species reported in clinical settings worldwide [13, 25]. Less frequently encountered taxa, including *S. aurantiacum*, *S. dehoogii*, and *S. ellipsoideum*, were also identified, in agreement with sporadic reports from Europe and the United States [13, 25]. Our dataset further included several rarely described species: *S. angustum*, *S. fusoideum*, *S. minutisporum*, *S. marinum*, *S. haikouense*, and *S. sphaerospermum*, which are seldom represented in susceptibility studies and often absent from large reference collections [11–13, 25]. These rare taxa exhibited distinctive antifungal profiles, and with the multidrug-resistant phenotype of *L. prolificans*, which remains resistant to all triazoles and amphotericin B but susceptible to olorofim [11–13, 15, 25].

We also identified *Scedosporium* sp. nov.1, a putative novel taxon not previously described. Phylogenetically distinct from known species, this isolate exhibited high triazole MICs that diverged from both *S. apiospermum* and *S. boydii* (**Table 1**, **Figure S3**), suggesting an emerging resistant lineage. This finding underscores the need for comprehensive molecular and phenotypic characterization of new taxa. Additionally, *S. multisporum* was detected among clinical isolates (**Table S1**), a species previously reported only from environmental sources [26]. This finding suggests that taxa traditionally considered environmental may also act as opportunistic pathogens. Overall, the detection of uncommon and novel species highlights the ecological diversity of *Scedosporium* and emphasizes the importance of accurate molecular identification, as rare or cryptic taxa may harbor distinct resistance phenotypes with direct clinical implications.

Clinically, our findings reaffirm the multidrug-resistant phenotype of *L. prolificans* and the limited therapeutic options available for *Scedosporium* infections. Current management relies heavily on surgical intervention and experimental antifungal combinations, yet outcomes remain poor [1, 2]. In this study, voriconazole retained activity against most *Scedosporium* species but was ineffective against *L. prolificans* (MICs ≥8 mg/L) (**Table 1**). In contrast, olorofim demonstrated potent *in vitro* activity against all *Scedosporium* and *Lomentospora* isolates, with MICs markedly lower than those of conventional antifungals (**Table 1**). These findings are consistent with multicenter studies reporting GM MICs <0.25 mg/L for both genera (**Table S2**). The current Taiwan dataset included not only clinical isolates but also the first large-scale collection of environmental reservoirs and novel taxa. These findings confirm the global consistency of olorofim susceptibility and show no evidence of regional resistance variation. Importantly, the uniformly low olorofim MICs observed here align with its favorable clinical responses reported in a recent Phase 2b trial [27].

Heatmap and correlation analyses demonstrated that antifungal susceptibility was primarily determined by species identity rather than isolate source (**Figures 3 and 4**). *S. apiospermum* and *S. boydii* displayed similar triazole activity patterns, whereas *L. prolificans* clustered separately, displaying pan-resistance to all agents except olorofim, consistent with its known multidrug-resistant profile [13, 25]. Several rare taxa, including *S. marinum*, *S. haikouense*, and *Scedosporium* sp. nov.1, exhibited divergent triazole MIC values but consistently low MICs to olorofim, although larger sample sizes are needed to confirm species-specific patterns. Correlation analyses revealed moderate positive associations among triazoles, consistent with their shared molecular target, while amphotericin B and olorofim demonstrated weak or no correlation with other agents, reflecting distinct mechanisms of action. Together, these findings reinforce the unique activity profile of olorofim across all tested species. Phylogenetic signal analysis (**Figure 5**) further demonstrated drug-dependent structuring, with olorofim activity conserved across lineages and triazole responses remaining heterogeneous, underscoring the value of integrating phylogenetic context into antifungal susceptibility assessments [10, 12, 13, 15, 25]. Given the variable and unpredictable triazole susceptibility among *Scedosporium* species, precise molecular identification remains essential, while olorofim emerges as the most promising therapeutic option for infections caused by these intrinsically resistant molds.

A key limitation of this study is the reliance on ITS and β-tubulin barcoding markers, which, although informative for taxonomy, do not encode antifungal targets and often fail to resolve cryptic species [28]. This limitation may partly explain the weak or inconsistent phylogenetic signal observed for triazoles and polyenes. Future investigations should incorporate target-associated loci such as *CYP51* and *DHODH* the molecular target of olorofim [10, 29, 30]. Phylogenies based on these genes could provide improved resolution and help clarify whether resistance traits in *Scedosporium* and *Lomentospora* represent phylogenetically conserved characteristics or have arisen through lineage-specific adaptation. Integrating genomic data with phenotypic susceptibility profiles will be essential to advance our understanding of resistance evolution in these emerging pathogens.

In conclusion, antifungal susceptibility in *Scedosporium* and *Lomentospora* was drug dependent. Olorofim exhibited uniformly low MICs and a conserved activity profile, whereas amphotericin B and triazoles displayed weak-to-moderate phylogenetic signals and marked heterogeneity, reflecting variable, species-specific responses. These findings highlight the promise of olorofim as a broadly active agent against intrinsically resistant molds and underscore the need for target-gene–based analyses to clarify the evolutionary basis of antifungal resistance.

## Acknowledgments

The pure compound olorofim was provided by F2G, Ltd.

## Transparency declaration

The authors declare no conflicts of interest. This study was supported by research grants from the Ministry of Science and Technology, Taiwan (NSTC 114-2311-B-037-002), and the Kaohsiung Medical University Research Program (KMU-TB114001).

## Author contributions

Lin SY and Huang YT conceived the study and contributed to data acquisition. Huang YT, Wu CJ, and Sun PL curated the data. Tseng YT, Abdrabo KAE, and Huang YT performed data analysis and interpretation. Lin SY, Huang YT, Sukphopetch P, and Chen SC drafted and edited the manuscript. Lin SY and Huang YT critically revised the manuscript. All authors reviewed and approved the final version of the manuscript.

## Notes

### Competing Interest Statement

The authors have declared no competing interest.

## References

1. Hoenigl M, Salmanton-Garcia J, Walsh TJ, Nucci M, Neoh CF, Jenks JD, et al. Global guideline for the diagnosis and management of rare mould infections: an initiative of the European Confederation of Medical Mycology in cooperation with the International Society for Human and Animal Mycology and the American Society for Microbiology. Lancet Infect Dis 2021;21:e246–e57.

2. Neoh CF, Chen SC, Lanternier F, Tio SY, Halliday CL, Kidd SE, et al. Scedosporiosis and lomentosporiosis: modern perspectives on these difficult-to-treat rare mold infections. Clin Microbiol Rev 2024;37:e0000423.

3. Marinelli T, Kim HY, Halliday CL, Garnham K, Bupha-Intr O, Dao A, et al. *Fusarium* species, *Scedosporium* species, and *Lomentospora prolificans*: A systematic review to inform the World Health Organization priority list of fungal pathogens. Med Mycol 2024;62:myad128.

4. Harun A, Gilgado F, Chen SC, Meyer W. Abundance of *Pseudallescheria*/*Scedosporium* species in the Australian urban environment suggests a possible source for scedosporiosis including the colonization of airways in cystic fibrosis. Med Mycol 2010;48 Suppl 1:S70–6.

5. Heath CH, Slavin MA, Sorrell TC, Handke R, Harun A, Phillips M, et al. Population-based surveillance for scedosporiosis in Australia: epidemiology, disease manifestations and emergence of *Scedosporium aurantiacum* infection. Clin Microbiol Infect 2009;15:689–93.

6. Rougeron A, Giraud S, Alastruey-Izquierdo A, Cano-Lira J, Rainer J, Mouhajir A, et al. Ecology of *Scedosporium* Species: Present Knowledge and Future Research. Mycopathologia 2018;183:185–200.

7. Wu HM, Fan YH, Phang GJ, Zeng WT, Abdrabo KAE, Wu YT, et al. Human activity, not environmental factors, drives *Scedosporium* and *Lomentospora* distribution in Taiwan. Med Mycol 2025;63:myaf022.

8. Cooley L, Spelman D, Thursky K, Slavin M. Infection with *Scedosporium apiospermum* and *S. prolificans*, Australia. Emerg Infect Dis 2007;13:1170–7.

9. Delhaes L, Harun A, Chen SC, Nguyen Q, Slavin M, Heath CH, et al. Molecular typing of Australian *Scedosporium* isolates showing genetic variability and numerous *S. aurantiacum*. Emerg Infect Dis 2008;14:282–90.

10. Vanbiervliet Y, Van Nieuwenhuyse T, Aerts R, Lagrou K, Spriet I, Maertens J. Review of the novel antifungal drug olorofim (F901318). BMC Infect Dis 2024;24:1256.

11. Halliday CL, Tay E, Green W, Law D, Lopez R, Faris S, et al. *In vitro* activity of olorofim against 507 filamentous fungi including antifungal drug-resistant strains at a tertiary laboratory in Australia: 2020-2023. J Antimicrob Chemother 2024;79:2611–21.

12. Escribano P, Gomez A, Reigadas E, Munoz P, Guinea J, Group AS. EUCAST-obtained olorofim MICs against *Aspergillus* and *Scedosporium* species and *Lomentospora prolificans* showed high agreements between visual inspection and spectrophotometric readings. Antimicrob Agents Chemother 2022;66:e0084922.

13. Rivero-Menendez O, Cuenca-Estrella M, Alastruey-Izquierdo A. *In vitro* activity of olorofim against clinical isolates of *Scedosporium* species and *Lomentospora prolificans* using EUCAST and CLSI methodologies. J Antimicrob Chemother 2020;75:3582–5.

14. Biswas C, Law D, Birch M, Halliday C, Sorrell TC, Rex J, et al. *In vitro* activity of the novel antifungal compound F901318 against Australian *Scedosporium* and *Lomentospora* fungi. Med Mycol 2018;56:1050–4.

15. Georgacopoulos O, Nunnally N, Law D, Birch M, Berkow EL, Lockhart SR. *In vitro* activity of the novel antifungal olorofim against *Scedosporium* and *Lomentospora prolificans*. Microbiol Spectr 2023;11:e0278922.

16. Risum M, Hare RK, Gertsen JB, Kristensen L, Rosenvinge FS, Sulim S, et al. Azole resistance in *Aspergillus fumigatus*. The first 2-year’s data from the Danish national surveillance study, 2018-2020. Mycoses 2022;65:419–28.

17. Tio SY, Chen SC, Hamilton K, Heath CH, Pradhan A, Morris AJ, et al. Invasive aspergillosis in adult patients in Australia and New Zealand: 2017-2020. Lancet Reg Health West Pac 2023;40:100888.

18. Steenwyk JL, Buida TJ, 3rd, Li Y, Shen XX, Rokas A. ClipKIT: A multiple sequence alignment trimming software for accurate phylogenomic inference. PLoS Biol 2020;18:e3001007.

19. CLSI. 2017. Reference method for broth dilution antifungal susceptibility testing of filamentous fungi, 3rd ed. CLSI document M38. Clinical and Laboratory Standards Institute, Wayne, PA.

20. Wickham H, François R, Henry L, Müller K, Vaughan D. dplyr 2023. https://dplyr.tidyverse.org (accessed May 6, 2025)

21. Wei T, Simko V, Levy M, Xie Y, Jin Y, Zemla J. Package “corrplot.” Statistician 2017.

22. Revell LJ. phytools: an R package for phylogenetic comparative biology (and other things): phytools: R package. Methods Ecol Evol 2012;3:217–23. 10.1111/j.2041-210X.2011.00169.x.

23. Oksanen J, Blanchet FG, Kindt R, Legendre P, Minchin PR, O’hara R, et al. Package ‘vegan’. Community ecology package, version 2. The Comprehensive R Network (CRAN) [Google Scholar] 2013.

24. Paradis E, Schliep K. ape 5.0: an environment for modern phylogenetics and evolutionary analyses in R. Bioinformatics 2019;35:526–8.

25. Czech MM, Cuellar-Rodriguez J, Kwon-Chung KJ, Stock F, Aneke CI, Olivier KN, et al. Clinical significance and antifungal susceptibility profile of 103 clinical isolates of *Scedosporium* species complex and *Lomentospora prolificans* obtained from NIH patients. J Clin Microbiol 2025;63:e0155024.

26. Zhang ZY, Shao QY, Li X, Chen WH, Liang JD, Han YF, et al. Culturable fungi from urban soils in China I: description of 10 new taxa. Microbiol Spectr 2021;9:e0086721.

27. Maertens JA, Thompson GR, 3rd, Spec A, Donovan FM, Hammond SP, Bruns AHW, et al. Olorofim for the treatment of invasive fungal diseases in patients with few or no therapeutic options: a single-arm, open-label, phase 2b study. Lancet Infect Dis 2025; doi: 10.1016/S1473-3099(25)00224-5.

28. Borman AM, Johnson EM. Name changes for fungi of medical importance, 2018 to 2019. J Clin Microbiol 2021;59;e01811–20.

29. Wu Y, Grossman N, Totten M, Memon W, Fitzgerald A, Ying C, et al. Antifungal susceptibility profiles and drug resistance mechanisms of clinical *Lomentospora prolificans* isolates. Antimicrob Agents Chemother 2020;64: e00318–20.

30. Celia-Sanchez BN, Mangum B, Brewer M, Momany M. Analysis of Cyp51 protein sequences shows 4 major *Cyp51* gene family groups across fungi. G3 (Bethesda). 2022;12:jkac249

